# AID-mediated protein knockdown reveals the requirement of NANOS2 in prenatal gonocytes for establishing functional spermatogonial stem cells

**DOI:** 10.1101/2025.05.22.655677

**Authors:** Yumiko Saga, Quan Wu

## Abstract

The RNA-binding protein NANOS2 plays a crucial role in male gonocyte development and the maintenance of spermatogonial stem cells. In the absence of the Nanos2 gene (Nanos2-KO), germ cells fail to enter G0 arrest and initiate the male differentiation program (including DNA methylation and the piRNA pathway), ultimately undergoing apoptosis before birth. *Nanos2* transcription begins at embryonic day 12.5 (E12.5) and terminates at E15.5. However, as the NANOS2 protein continues to be stably expressed beyond E15.5, it is important to elucidate the function of NANOS2 during this post-E15.5 period in germ cell fate determination. To address the functional significance of sustained NANOS2 protein expression, we employed an auxin-inducible degron (AID2) system to achieve rapid degradation of NANOS2 after E15.5. Within 24 hours of 5-Ph-IAA administration, NANOS2 protein was efficiently depleted. As a result, germ cells resumed the cell cycle, exhibited aberrant gene expression patterns similar to Nanos2- KO gonocytes, and underwent apoptosis if NANOS2 depletion occurred at E15.5 or E16.5. Although some surviving cells initiated spermatogenesis and expressed PLZF and GFRA1 after birth, further spermatogenesis was not observed. These findings reveal that sustained NANOS2 protein expression during the embryonic stage is essential for establishing functional spermatogonial stem cells, highlighting a previously unrecognized regulatory mechanism in male germ cell development.

## Introduction

Spermatogenesis is a highly orchestrated process that ensures the continuous production of sperm throughout the male reproductive lifespan(De Rooij, 2017). The foundation for this process is established during embryogenesis when male germ cells, known as gonocytes, undergo a series of differentiation steps to form spermatogonial stem cells (SSCs)(Law and Oatley, 2020). These SSCs serve as the self-renewing progenitor population that maintains spermatogenesis throughout adulthood. Among the various factors implicated in this process, NANOS2, an RNA-binding protein, has been identified as a critical determinant of male germ cell fate. NANOS2 is expressed in embryonic male germ cells after E12.5 to play essential roles in establishing male properties, such as cell cycle arrest and DNA methylation during the embryonic stage(Suzuki and Saga, 2008). Mechanistically, NANOS2 makes a ternary complex with another RNA-binding protein DND1, and CNOT1, a component of the deadenylation complex, to repress the expression of target RNAs via either RNA degradation or translational repression(Hirano et al., 2022). However, in the absence of NANOS2, many male-specific genes, such as *Dnmt3l, Tdrd1*, and *Miwi2,* which are involved in DNA methylation, fail to be induced (Suzuki et al., 2010). Therefore, NANOS2 is a male-promoting factor, although the precise mechanism is unknown. After birth, NANOS2 is expressed in spermatogonial stem cells to maintain the stemness during spermatogenesis via repressing the mTORC1 signaling pathway(Zhou et al., 2015). Those functional analyses were conducted using gene-knockout mouse lines, either conventional or conditional *Nanos2*-knockout (KO). However, an important limitation of these studies is that protein expression does not always correlate with transcriptional activity. For example, *Nanos2* transcription starts at E12.5 and ceases at E15.5(Pui and Saga, 2017; Shimada et al., 2021), but the protein expression is maintained throughout the embryonic stage and persists even after birth. Prenatal expression of NANOS2 might be required to preserve gonocytes or initiate spermatogenesis. It is also possible that it is needed to establish spermatogonial stem cells, given its essential role in maintaining stemness(Sada et al., 2009). However, gene-KO technology is useless in testing these possibilities because NANOS2 protein expression continues after transcription is terminated at E15.5. To circumvent this limitation, we employed the auxin-inducible degron (AID) system, where a target protein is fused to a short peptide tag (AID-tag)(Nishimura et al., 2009). When the plant hormone auxin or the analogue is added, it recruits an F-box protein (such as *TIR1(F74G))* that leads to ubiquitination and rapid proteasomal degradation of the targeted protein(Yesbolatova et al., 2020). By leveraging this approach, we could specifically deplete NANOS2 protein after E15.5, allowing us to dissect its role in male germ cell development and SSC establishment in a precise temporal manner.

## Results

### 1. Establishment of AID-tagged NANOS2 knock-in mouse

A Cas9-mediated knockin method was employed to introduce an AID-tag with 3xFLAG at the N-terminal of the *Nanos2* coding sequence (Fig. S1). PCR screening identified multiple recombinant mice. Although most recombinants displayed the expected bands for 3′ recombination, the 5′ band sizes varied, suggesting insertions and/or deletions and mosaicism in F0 mice (Fig. S1). Sequence analysis revealed that none of the recombinants exhibited the expected recombination event. In most cases, deletions and insertions occurred at the Cas9 target site. However, one recombinant (#15) was found to have an in-frame insertion of the 3×FLAG and AID sequences at the N-terminus, along with an additional 24 N-terminal amino acids (Fig. S1). This male remained fertile even in the homozygous condition (*Nanos2^A/A^*). Since NANOS2-null males lack sperm, we expected this line (#15) to produce functional NANOS2 with the 3×FLAG and AID tag at the N-terminus. To confirm protein expression and degradation via the AID2 system, we crossed this line with the previously established *Oct-dPE-TIR1(F74G)-FLAG* mouse line (Fig. S2A). In wild-type embryos, NANOS2 expression remains stable from E16.5 to E18.5 (Fig. S2B), even after transcription is terminated at E15.5. A *Nanos2^A/A^* female was mated with a *Nanos2^A/+^/Oct-dPE-TIR1(F74G)-FLAG* male, and the resulting embryos were injected with 5-Ph-IAA for three consecutive days starting at E13.5. Testes were collected from embryos at E16.5 and analyzed by western blotting and immunohistology. Anti-FLAG antibodies detected FLAG-tagged NANOS2 and TIR1(F74G). The results showed that NANOS2 protein was successfully degraded in the presence of TIR1(F74G), although some residual NANOS2 protein was observed in homozygous *Nanos2^A/A^* testes (Fig. S2C). Histological analysis supported this finding: E-cadherin (CDH), a marker of embryonic germ cells, co-localized with NANOS2 in *Nanos2^A/A^* testes, whereas NANOS2 signals disappeared in the presence of TIR1 (Fig. S2D), confirming efficient protein knockdown via the AID2 method.

### 2. NANOS2 protein was eliminated within 24 hours in vivo

Having established that NANOS2 could be efficiently depleted using the AID system, we next investigated the consequences of its loss. Once male germ cells express NANOS2 after E13.5, they stop proliferation and enter the G0 arrest stage. The DNA methylation program is initiated by the expression of DNMT3L after E14.5. Therefore, male gonocyte properties may be established by E15.5. *Nanos2* transcription stops at E15.5 but NANOS2 protein is maintained until birth (Fig. S2B). Thus, experiments were designed to knock down NANOS2 protein after E15.5 by injecting 5-Ph-IAA at E15.5, 16.5, 17.5, and 18.5 (Fig. 1A). We established *Nanos2^A/A^*line with and without *Oct4-dPE- TIR1(F74G)-Flag*. Thus, the genotype of analyzed embryos was *NANOS2^A/A^/Oct-dPE- TIR1* or *NANOS2^A/A^* (serve as control).

**Figure 1.**
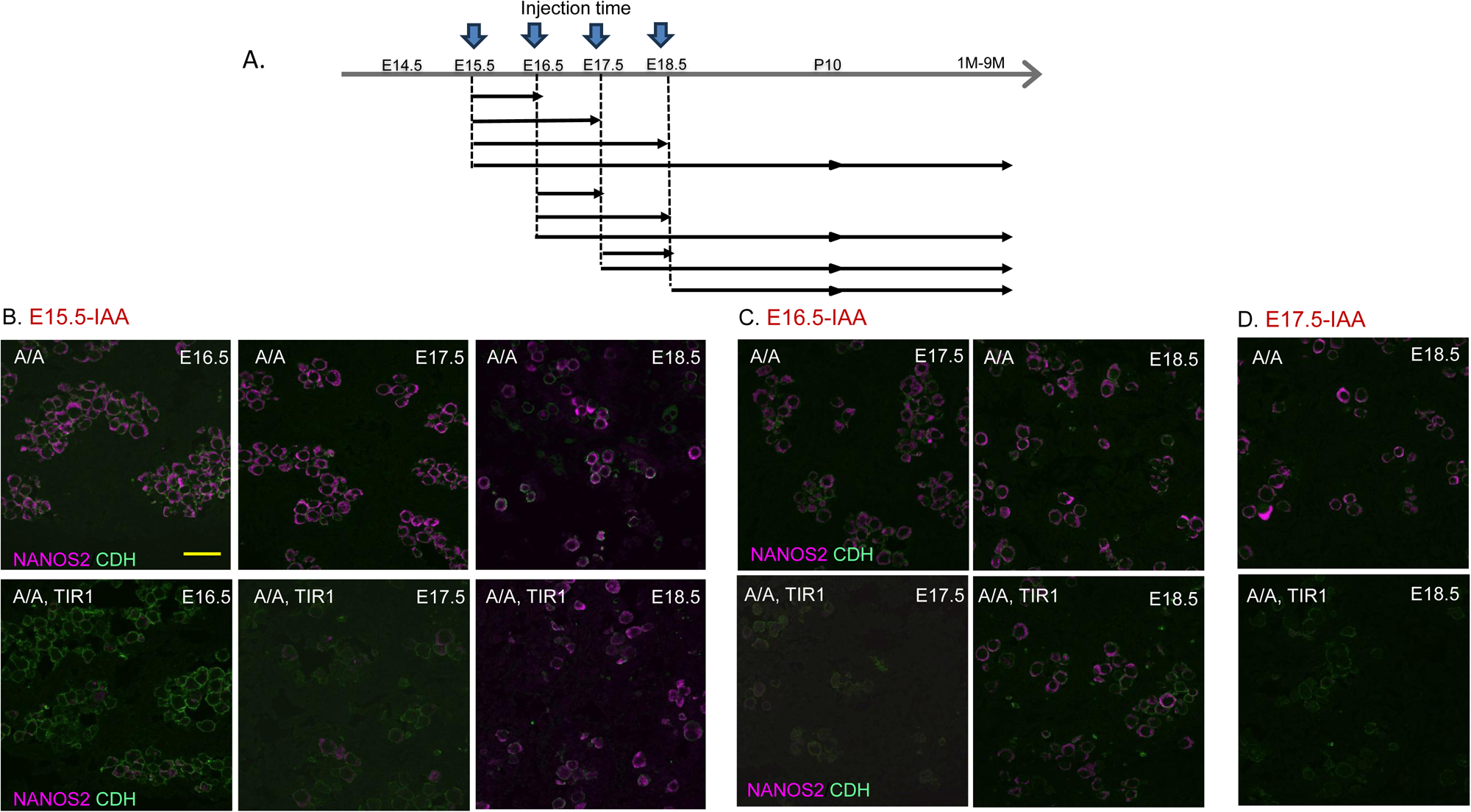
Successful NANOS2 knockdown via AID-mediated method. (A) Experimental scheme. Pregnant females were subjected to single or consecutive injections of 5-Ph-IAA between E15.5 and E18.5. Embryonic testes were prepared 24, 48, and 72 hours after injection up to E18.5. Testis samples were also prepared after birth on D10 and 1-6 months. (B) Immunofluorescent signals of NANOS2 (magenta) and CDH (green). Injection times are indicated by red letters. Each genotype and sampling time is indicated in each panel. (A/A) indicates homozygous *AID-Nanos2*. Scale bar, 30 μm.

First, we examined NANOS2 expression by immunohistochemistry using an anti- NANOS2 antibody. When 5-Ph-IAA was injected at E15.5 (E15.5-IAA), NANOS2 protein was already eliminated by E16.5, and this condition was maintained until E17.5 (Fig. 1B). However, NANOS2 protein was detected again at E18.5 (Fig. 1B). A similar result was observed when 5-Ph-IAA was injected at E16.5; NANOS2 protein was eliminated by E17.5 but reappeared at E18.5 (Fig. 1C). In contrast, when IAA was administered at E17.5, NANOS2 protein was not detected at E18.5 (Fig. 1D). Since Nanos2 transcription terminates at E15.5, this protein recovery was unexpected. The most likely explanation is the reinitiation of transcription. To investigate this possibility, we used RNAscope (biotech) to detect *Nanos2* RNA. As expected, strong *Nanos2* signals were detected at E14.5 but disappeared by E15.5 in wild-type embryos, confirming the cessation of transcription. While no signals were detected in the E18.5 control sample, a few signals were observed in NANOS2-positive cells of the E15.5-IAA sample, indicating that protein expression resulted from newly synthesized transcripts (Fig. S3).

### 3. Recapitulation of NANOS2 loss phenotype

To understand the molecular changes associated with NANOS2 depletion, we examined the expression of key germ cell regulatory factors. One of the most obvious phenotypes in *Nanos2*-null germ cells is the loss of DNMT3L expression, a cofactor of DNA methyl transferase DNMT2a/2b required for establishing male-type epigenetic status (Kato et al., 2007). DNMT3L expression begins at E14.5 following NANOS2 expression in the wild-type but is not induced without NANOS2. We did not expect any abnormality when NANOS2 protein was knocked down at E15.5 (E15.5-IAA) since DNMT3L expression would have already been initiated. As expected, DNMT3L expression was detected in IAA-treated gonocytes. However, we observed lower expression in E15.5-IAA gonocytes (Fig. 2A). Interestingly, DNMT3L expression was unchanged of E16.5-IAA case, indicating that NANOS2 expression is required until E16.5 to achieve stable DNMT3L expression. Then we checked another male germ cell property, cell cycle status by Ki67 immunostaining (Fig. 2B). Previous studies have shown that NANOS2 is involved in both the entry and maintenance of the G0 phase of the cell cycle (Shimada et al., 2021). As expected, Ki67 signals were detected in E15.5-IAA gonocytes at E17.5 but not at E16.5 (Fig. 2B). A similar pattern was observed in E16.5-IAA gonocytes, where Ki67 signals were absent at E17.5 but reappeared at E18.5 (Fig. 2B). These results indicate that continuous NANOS2 expression is required to maintain the G0 state in male gonocytes. Another well-known feature of the NANOS2 loss-of-function phenotype is the upregulation of NANOS3, which partially compensates for the function of NANOS2. Similar to Nanos2 knockout (KO) mice, NANOS3 expression was upregulated in E15.5- IAA gonocytes, even at E16.5 (Fig. S4A). NANOS3 functions as an anti-apoptotic factor in primordial germ cells (PGCs)(Tsuda et al., 2003) and gonocytes(Suzuki et al., 2014). However, despite the upregulation of NANOS3, apoptotic cell death was observed in IAA-treated gonocytes (Fig. S4B). This phenotype is also observed in NANOS2-null germ cells(Suzuki and Saga, 2008).

**Figure 2.**
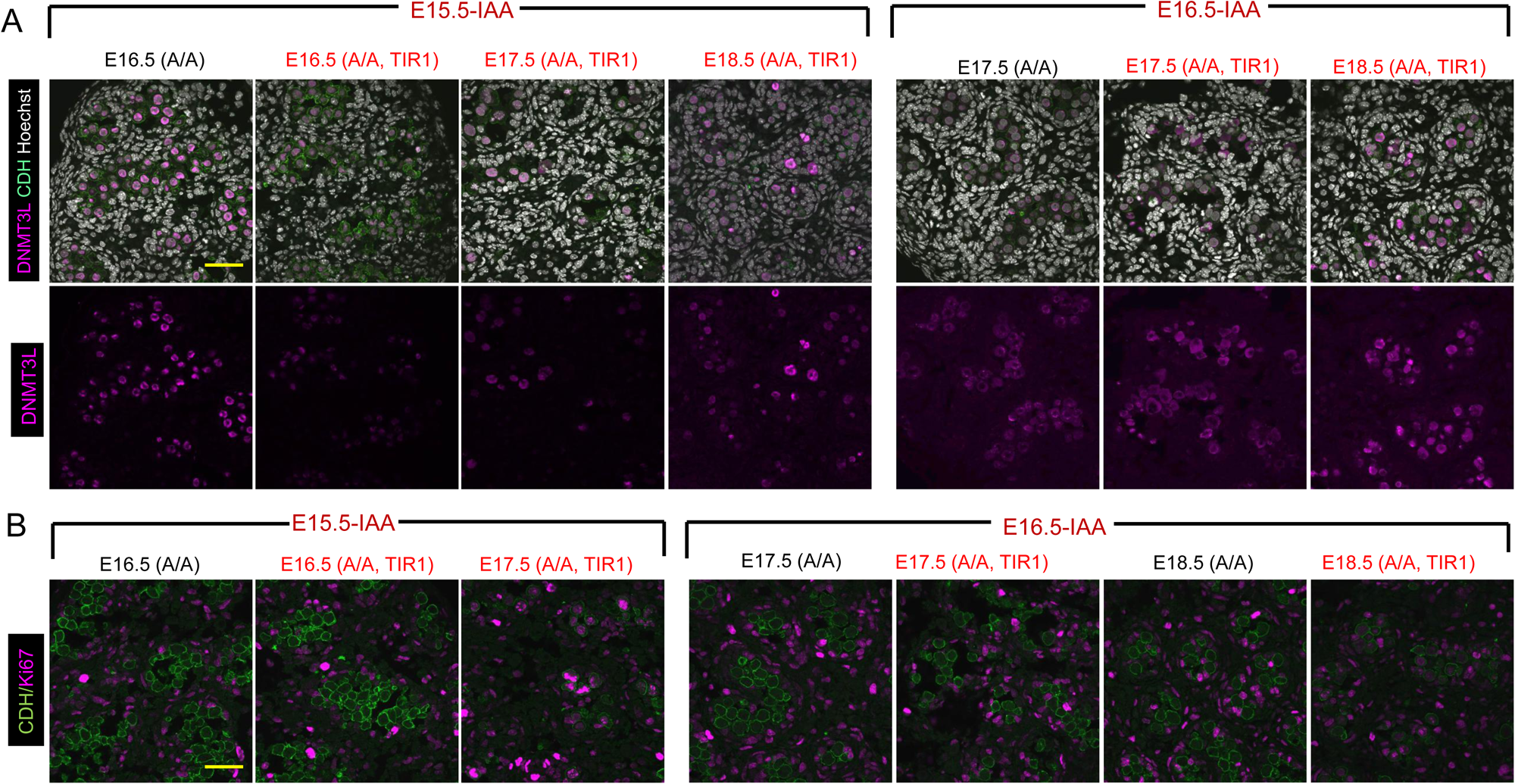
Impact of NANOS2 knockdown during embryonic stages. (A) Immunofluorescent images of DNMT3L (magenta) and CDH (green) expression after NANOS2 knockdown at E15.5 and E16.5. (B) Immunofluorescent signals of Ki67 (magenta) and CDH (green) after NANOS2 knockdown at E15.5 and E16.5. Scale bar, 30 μm.

Another intriguing feature of NANOS2-null germ cells is the premature expression of PLZF, a marker of undifferentiated spermatogonia (Costoya et al., 2004). PLZF expression is detectable during embryonic development, but its intensity increases significantly after birth. However, in E15.5-IAA gonocytes, PLZF expression was already enhanced, particularly at E18.5 (Fig. S4C). In general, the absence of NANOS2 prevents male-type differentiation. However, based on the PLZF expression pattern, the loss of NANOS2 may induce the premature differentiation of postnatal spermatogonia. This feature has also been reported in *Nanos2-*KO gonocytes at E16.5(Pui and Saga, 2018).

### 4. Contribution of embryonic NANOS2 to spermatogenesis

We next examined the long-term effects of NANOS2 depletion on germ cell survival and SSC establishment. The elimination of NANOS2 after E15.5 induced phenotypes similar to those observed in NANOS2-null germ cells, which include apoptotic cell death(Figure S4B). However, some germ cells survived and expressed PLZF. To ask if those gonocytes retain the ability to differentiate into spermatocyte-and if so, how long NANOS2 expression during the embryonic stage is required for spermatogenic differentiation-we examined male progenies treated by 5Ph-IAA at E15.5, E16.5, E17.5, and E18.5, 9-10 days after birth. In wild-type mice, male spermatogonia resume cell proliferation 1.5 days after birth. As a result, seminiferous tubules in control testes are densely packed with proliferating spermatogonia (Fig. 3). In contrast, many empty tubules were observed in E15.5-IAA and E16.5-IAA testes. This phenotype became milder in E17.5-IAA testes, and no obvious changes were visible in E18.5-IAA testes (Fig. 3, upper panel). Spermatogonial cells can be classified into differentiating and undifferentiated spermatogonia(Yoshida et al., 2006). PLZF is a marker for undifferentiated spermatogonia (Costoya et al., 2004). Although the number of PLZF-positive cells was lower in E16.5-IAA testes, a considerable number were still detected (Fig. 3, center panel). Similarly, NANOS2 expression was observed in all conditions (Fig. 3, bottom panel), indicating that the transient elimination of NANOS2 after E15.5 did not prevent the formation of undifferentiated spermatogonia. PLZF is not a definitive stem cell marker, but spermatogonial stem cells, which are characterized by the expression of both NANOS2 and GFRA1 may emerge from the PLZF-positive cell population. If stem cells are generated, normal spermatogenesis should be observed in IAA-treated testes. To address this issue, we examined IAA-treated males at later stages. In 2.5-month-old (10- week-old) control testes, many tubules were filled with spermatogonia and differentiating spermatocytes, including elongated spermatids (Fig. 4A). Similarly, relatively normal tubules were observed in both E17.5-IAA and E18.5-IAA testes at 3.5 months. In contrast, E16.5-IAA testes examined at 2.5 months contained many empty tubules. Nevertheless, PLZF-positive cells were found at the periphery of seminiferous tubules, even in severely affected E16.5-IAA tubules. Additionally, GFRA1-positive cells were detected among the PLZF-positive cells, indicating the presence of spermatogonial stem cells. Furthermore, comparable levels of GFRA1-positive cells were observed in E16.5-IAA testes compared with control, E17.5-IAA, and E18.5-IAA testes (Fig. 4A, Table S1). These results suggest that continuous expression of NANOS2 after E15.5 may not be strictly required to initiate spermatogenesis or establish spermatogonial stem cells. However, NANOS2 knockdown may not have been complete, and some germ cells that escaped NANOS2 depletion could have contributed to the production of spermatogenic cells. To rule out this possibility, we used *Nanos2(A/mch)* mice, which may express a lower level of NANOS2 protein, making escape less likely. We treated these mice with 5-Ph-IAA twice, at E15.5 and E16.5, and examined the progeny after birth. At one month, we observed many PLZF-positive cells and some PLZF/GFRA1-double-positive cells, even though spermatogenesis was severely compromised, indicating the presence of spermatogenical stem cells (Fig. 4B, Table S1). However, normal spermatogenesis never recovered in either E16.5-IAA or E15.5+E16.5-IAA testes, even after six months (Fig. 4C, Fig. 5). We also examined 9-12- month-old testes. Still, testis size was petite (Fig. 5). In E17.5-IAA mice, testis size was variable. In contrast, normal-sized testes were recovered in E18.5-IAA mice (Fig. 5). These results indicate that continuous expression of NANOS2 during the embryonic stage is essential for initiating normal spermatogenesis and producing functional spermatogonial stem cells.

**Figure 3.**
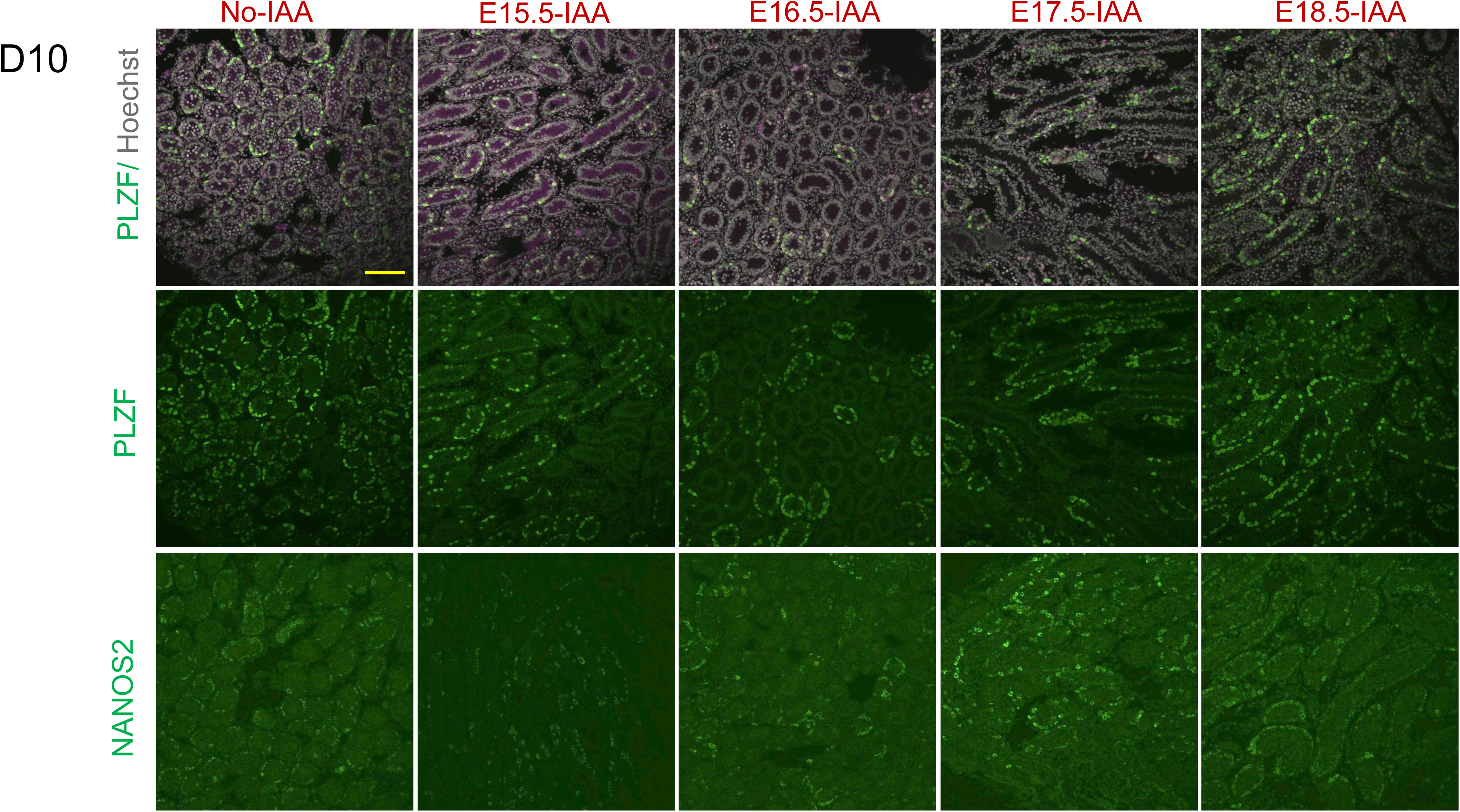
Impact of NANOS2 knockdown for initiation of spermatogenesis. Immunofluorescent signals of PLZF (with and without Hoechst) and NANOS2 at postnatal day 10. Scale bar, 100 μm.

**Figure 4.**
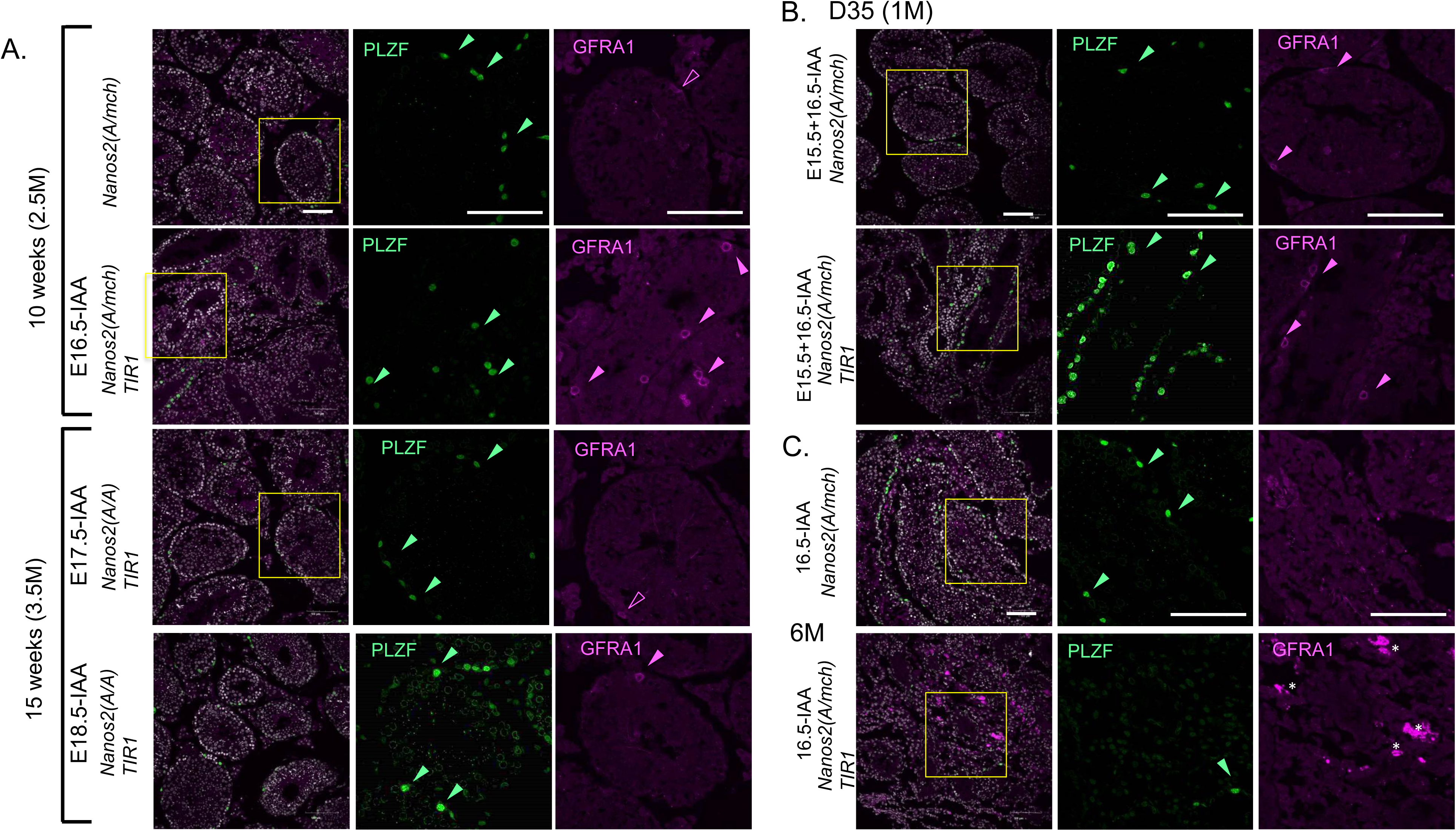
Impact of NANOS2 knockdown during spermatogenesis. Immunofluorescent signals of PLZF and GFRA1 at 10 weeks (A), D35(B), and 6M (C) after birth. Enlarged images of square regions are shown as single channels for PLZF and GFRA1. Arrowheads indicate representative signals. Asterisks are non-specific signals strongly observed in the 6M NANOS2-knockdown sample. *Nanos2 (A/mch)* means double heterozygous of *AID- Nanos2* and *Nanos2-mCherry* (KO) alleles. Scale bar, 100 μm.

**Figure 5.**
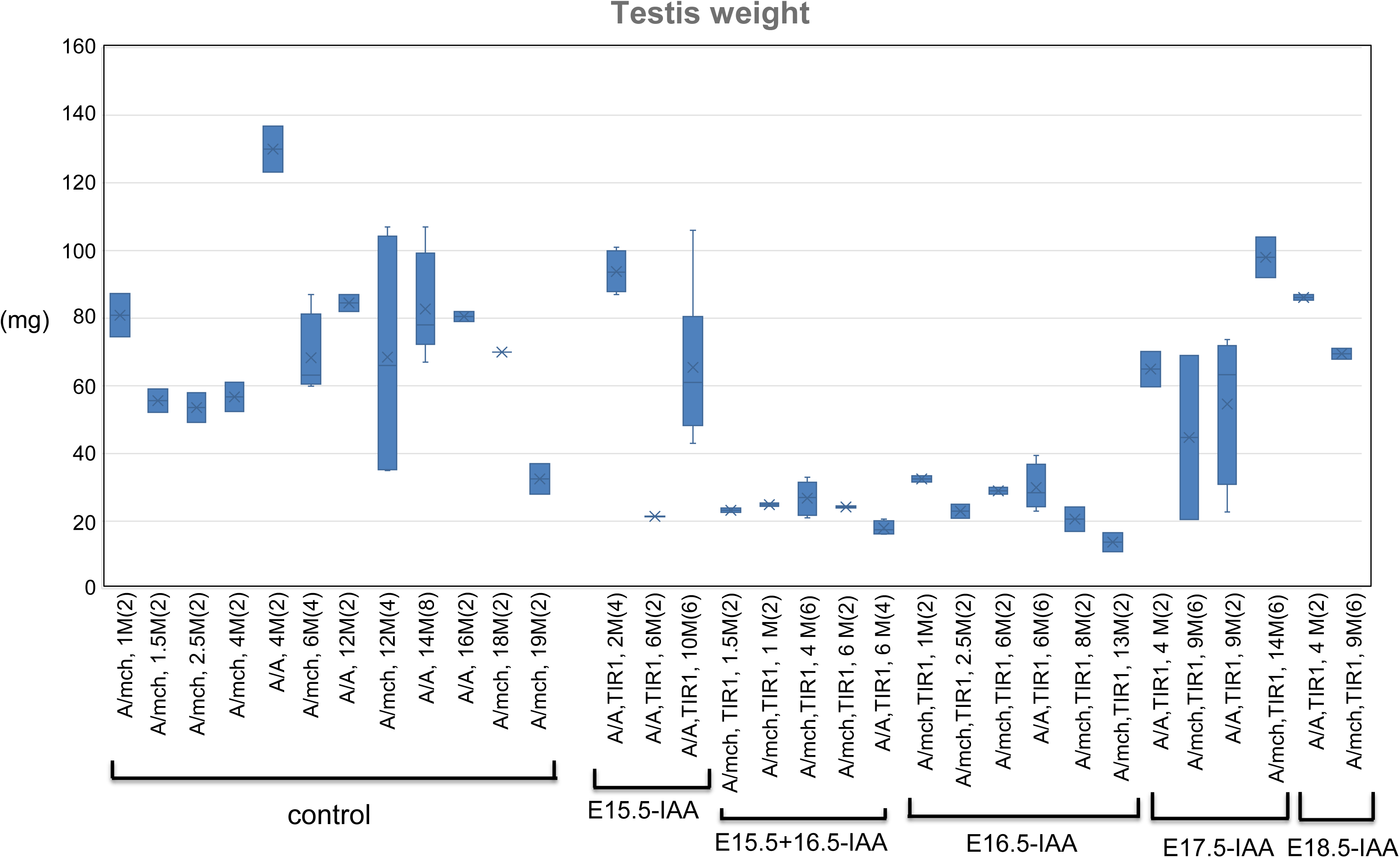
Quantification of testis weight for control mice without TIR1 (Nanos2A/A or Nanos2A/mch) and IAA-treated mice at various embryonic stages (E15.5, E15.5+E16.5, E16.5, E17.5, and E18.5). The age of the sacrificed mice and the number (in parentheses) of testes measured are indicated after the genotype in each case.

## Discussion

Direct regulation of proteins has been challenging due to the lack of effective methods. We applied AID technology to investigate the role of NANOS2 after transcription ceases during the perinatal stage. NANOS2 transcription ceases at E15.5, by which point key genetic events leading to male gonocyte differentiation may be completed. For instance, by E15.5, all gonocytes halt proliferation and enter the G0 phase(Shimada et al., 2021). Additionally, numerous factors involved in epigenetic modifications, such as DNA methylation and the PiRNA pathway, are expressed by E15.5 (Suzuki et al., 2010). Therefore, we initially expected that NANOS2 protein observed beyond this stage is not essential for germ cell development. However, our protein knockdown experiment clearly revealed that continuous NANOS2 protein expression is critical to establish spermatogonial stem cells.

We performed protein knockdown at different time points after E15.5 and observed distinct effects on DNMT3L expression between E15.5-IAA and E16.5-IAA conditions.

Specifically, DNMT3L expression was downregulated when NANOS2 was removed at E15.5 (E15.5-IAA), whereas the effect was minimal in E16.5-IAA. This indicates the existence of a critical window in which stable DNMT3L expression depends on NANOS2 function. The molecular mechanism underlying DNMT3L activation remains unclear. Although NANOS2 is not a transcription factor but rather an RNA-binding protein that represses target RNAs, many genes are activated depending on NANOS2 function. Previous studies have shown that ectopic NANOS2 expression in female germ cells leads to the activation of several male-specific genes, including DNMT3 (A. Suzuki & Saga, 2008). This suggests that NANOS2 alters the epigenetic landscape to favor male differentiation. Since DNMT3L expression was maintained in E16.5-IAA, likely, a male-specific epigenetic program is already established by this stage. Nevertheless, the E16.5-IAA group failed to undergo normal spermatogenesis after birth (Figure S5). In contrast, E17.5-IAA exhibited a milder phenotype, and E18.5-IAA showed no defects in spermatogenesis, indicating that the transient loss of NANOS2 around E16.5 was detrimental to subsequent spermatogenesis. Intriguingly, PLZF/GFRA1 double-positive cells were frequently observed in severely affected testes, such as those in E16.5-IAA and E15.5+E16.5-IAA groups, at 1–3 months of age. If these cells functioned as spermatogonial stem cells, spermatogenesis would be resumed later. However, spermatogenesis never recovered, even after more than 10 months (Figure S5), suggesting that the PLZF/GFRA1 double-positive cells observed in E16.5-IAA testes at 1–3 months were defective as stem cells. These findings appear to contradict previous reports from transplantation experiments, which demonstrated that spermatogonial stem cells could be derived from embryonic germ cells both before (E6.5–E12.5) and after (E14.5–E16.5) NANOS2 expression (Chuma et al., 2005; Ohta et al., 2004). Thus, the strong impact of transient NANOS2 loss was unexpected and warrants further investigation. We observed the reappearance of NANOS2 protein at E18.5 in both cases of E15.5-IAA and E16.5- IAA due to transcriptional activation, indicating a negative feedback mechanism of *Nanos2* transcription by the product. Nevertheless, normal spermatogenesis was not rescued by the induced NANOS2, indicating again the importance of continuous expression at a critical time window (E15.5-E17.5). We hypothesize that an irreversible epigenetic change induced by the absence of NANOS2 at this critical window may contribute to fatal defects in germ cells. The AID2-mediated knockdown system provides an ideal approach to directly assess the immediate effects of NANOS2 loss and its role in the production of functional spermatogonial stem cells.

## Materials and Methods

### Animals

All animals were kept in a room conditioned at 23 ± 2 °C, with 50 ± 10% humidity, under a 12-hour light-and-dark cycle. All protocols and procedures involving the care and use of animals were reviewed and approved by the Institutional Animal Care and Use Committee of the National Institute of Genetics. Throughout the study, the care and use of animals were conducted under the guidelines and regulations set by the Ministry of Education, Culture, Sports, Science and Technology, the Ministry of the Environment, and the Science Council of Japan. The mouse line used for the production (B6/C3H-F1), and maintenance (ICR) were purchased from CLEA Japan, Inc. The production and characterization of *Oct-dPE-TIR1 (F74G)* transgenic mouse line have been described before(Makino-Itou et al., 2024). dNanos2-mCherry KO/KI mouse line was also described before(Wright et al., 2021).

### Generation of AID-Nanos2 knock-in mouse

A targeting vector was constructed in the *pBluescript-KS* vector, which includes *Nanos2 5’-UTR* (745 bp), *3X Flag-tag, AID-tag* with *Nanos2*-coding sequence, and the *3’ UTR* (472 bp). To induce homologous recombination using CAS9-mediated technology, the single-stranded targeting vector was generated using the TAKARA Guide-it™ Long ssDNA Production System (632644). The single-stranded DNA was injected with sgRNA (IDT) and TrueCut Cas9 protein v2 (Invitrogen) into the pronucleus of fertilized eggs (B6C3F1). F0 mouse tail DNA was used for PCR to detect homologous recombination events at 5’ and 3’ integration sites.

### Genetic cross and sample preparation

For sample preparation, homozygous *AID-Nanos2* (marked as *A/A*) male and female with or without *Oct-dPE-TIR1(F74G)-FLAG* (marked as *TIR1*) lines were used to set up mating of either *Nanos2^A/A^, TIR1* male and *Nanos2^A/A^* female or *Nanos2^A/A^* male and *Nanos2^A/A^*, TIR1 female to obtain *Nanos2^A/A^*, TIR1 and *Nanos2^A/A^*(control) embryos or pups. We occasionally crossed with NANOS2-null (*Nanos2-mcherry*) mouse line established before(Wright et al., 2021) to reduce *Nanos2* dosage. Homozygous NANOS2- null (*Nanos2^mch/mch^*) male is sterile, but a double heterozygous mouse *Nanos2^mch/A^* male is fertile. The date of the plug was denoted as E0.5. To induce protein degradation, 5-Ph-IAA (BioAkademia, Japan, #30-003) dissolved in PBS (0.5 mg/ml) was intraperitoneally injected (final concentration is 5 mg/kg) at the indicated time. Testis samples were fixed with 4% paraformaldehyde, embedded with OCT compound (Tissue Tek, Sakura) after sequential sucrose treatment (10% to 30%), and frozen.

### Western blotting

Testis samples were immediately frozen in liquid nitrogen. Frozen samples were lysed in TNE buffer (50 mM Tris-HCl pH 7.4, 150 mM NaCl, 1 mM DTT, 1 mM EDTA, and 1% NP40) with cOmplete Protease inhibitor cocktail (Roche). Appropriate amounts were mixed with 2x SDS sample buffer (Tris-HCl pH 6.8, 4% SDS, 20% glycerol, 10% 2- mercaptoethanol, and 0.004% bromophenol blue) and incubated at 95°C for 5 min before loading. After electrophoresis, proteins were transferred onto an Immobilon-P Transfer Membrane (Millipore). The membrane was incubated with a primary antibody in skim milk at 4°C overnight and subsequently incubated with a secondary antibody at room temperature for 2-3 hours. Detection was performed using the SuperSignal West Femto Maximum Sensitivity Substrate (Thermo Scientific), and images were acquired with a ChemiDoc Touch MP system (Bio-Rad). To detect Flag-tagged AID-NANOS2 and TIRI(F74G), anti-FLAG-M2-HRP (Sigma-Aldrich, A8592) was used at a 1:5000 dilution.

### Immunohistochemistry

Frozen sections prepared at 6 μm thickness were incubated with 3% BSA for 1 hour and subjected to the primary antibodies overnight at 4°C. After washing with PBST (PBS containing 0.1% Tween), sections were incubated with secondary antibodies containing bisbenzimide H33342 for 2 hours at RT. After washing with PBST, the sections were mounted and observed with an Olympus FV1200 or FV3000 confocal microscope. Primary antibodies were used at the following dilutions: goat anti-E-cadherin (1:400, R&D Systems, AF748), rabbit anti-NANOS2 (1:500, (Suzuki and Saga, 2008)), rabbit anti-DNMT3L (1:300, gift from Dr. Shinya Yamanaka), rabbit anti-NANOS3 (1:500, (Suzuki et al., 2009)), rabbit anti-Ki67 (1:200, Invitrogen, Cat # MA5-14520), rabbit anti-PLZF (1:200, Santa Cruz Biotechnology, sc-22839) and rabbit anti-Cleaved Caspase (1:200, Cell Signaling, D174, 5A1E, #9664). For secondary antibodies, donkey anti-rabbit and anti-goat antibodies conjugated with either Alexa-594 or Alexa-647 (Invitrogen) were used at a dilution of 1:1000.

### RNAscope *in situ* hybridization

For in situ hybridization, gonads were fixed in 4% paraformaldehyde, followed by sequential sucrose treatment (10% to 30%), embedded in OCT compound (Tissue-Tek, Sakura), and frozen. Sections were cut at 12 µm thickness. In situ hybridization was performed using the RNAscope™ Multiplex Fluorescent Reagent Kit v2 (#323100) according to the manufacturer’s instructions, with a probe targeting *Nanos2*. Following hybridization, sections were blocked in 3% skim milk in PBST (0.1% Tween-20 in PBS) for 1 hour at room temperature. Primary antibodies, goat anti-E-cadherin (1:400) and rabbit anti-NANOS2 (1:400), both diluted in 3% milk/PBST, were applied and incubated overnight at 4°C. After washing, secondary antibodies (Invitrogen, 1:1000 in PBST) were added for 1 hour at room temperature. Images were acquired using a Nikon A1 confocal microscope.

## Acknowledgements

We thank Ms. Noriko Yamatani for producing the AID-tagged *Nanos2* mouse line and Dr. Akemi Okubo, Ms. Yuko Katayama, and Hisako Inoue for supporting mouse care and experiments. We also acknowledge members of the Brain Function Laboratory (NIG) for providing experimental space and reagents. This work was supported by JSPS KAKENHI Grant Number 17H06166 and an AMED NBRP Fundamental Technologies Upgrading Program to Y. S.

## Author contributions

Y. S. designed the experiments and conducted most of them. Q.W. performed the *in situ* hybridization experiment. Y.S. and Q.W. wrote the manuscript.

## Supplementary Figure legends

Figure S1. Targeting strategy to generate AID-tagged NANOS2 and the detail of AID- NANOS2 allele. The targeting vector was designed to add 3xFLAG and AID tag at the end of NANOS2. The established line contained an additional N-terminal (24 amino acids) at the end of the recombinant. The detailed sequence data are shown.

Figure S2. Elimination of the NANOS2 protein via AID-mediated protein knockdown.

(A) Immunofluorescent signal of NANOS2 protein at E16.5-E18.5 testis sections. Scale bar, 30 μm. (B) Schematic presentation of AID-Nanos2 (A) allele and transgene expressing TIR1(F74G) only germ cells under the control of Oct4-delta PE promoter and enhancer (ref). (C) Western blot analysis of E16.5 testes prepared from pregnant females who received daily injections of 5-Ph-IAA from E13.5 to E15.5. Each genotype is indicated upper site of the blot. A/A mean homozygous AID-Nanos2 allele. TIR1 mean integration of *Oct-dPE-TIR1(F74G)* transgene. We used an anti-FLAG antibody to detect 3xFLAG-AID-tagged NANOS2 and FLAG-tagged TIR1(F74G). There are two specific bands for TIR1-FLAG, which could be derived from two possible in-frame translation start sites. (D) Immunofluorescent signals of germ cell marker. E-cadherin (CDH) and NANOS2. Blue signals are counter-staining with Hoechst. Scale bar, 50 μm.

Figure S3. NANOS2-knockdown may induce *Nanos2* transcription.

Co-staining of *Nanos2* transcripts and NANOS2 proteins at the E14.5, E15.5, and E18.5 wild type testes and E18.5 testis of E16.5-IAA. Arrowheads indicate *Nanos2* mRNA signals detected in NANOS2-positive cells of E16.5-IAA at E18.5. No such signal was detected in the wild-type control at E18.5.

Figure S4. Impact of NANOS2 knockdown during embryonic stage. (A) Immunofluorescent signals of NANOS3 and CDH. Scale bar, 20 μm. (B) Cleaved caspase signals indicating apoptotic cell death were frequently observed in sections of Nanos2(A/A, TIR1). Scale bar, 100 μm.

Figure S5. Summary of immunofluorescent data. Results were shown as comparative marks (- and +) to the control samples (A/A without TIR1). In the panel, only the E18.5 result of E15.5-IAA was shown as a control.

