## supplementary figs for "AID-mediated protein knockdown reveals the requirement of NANOS2 in prenatal gonocytes for establishing functional spermatogonial stem cells"

### Fig. S1

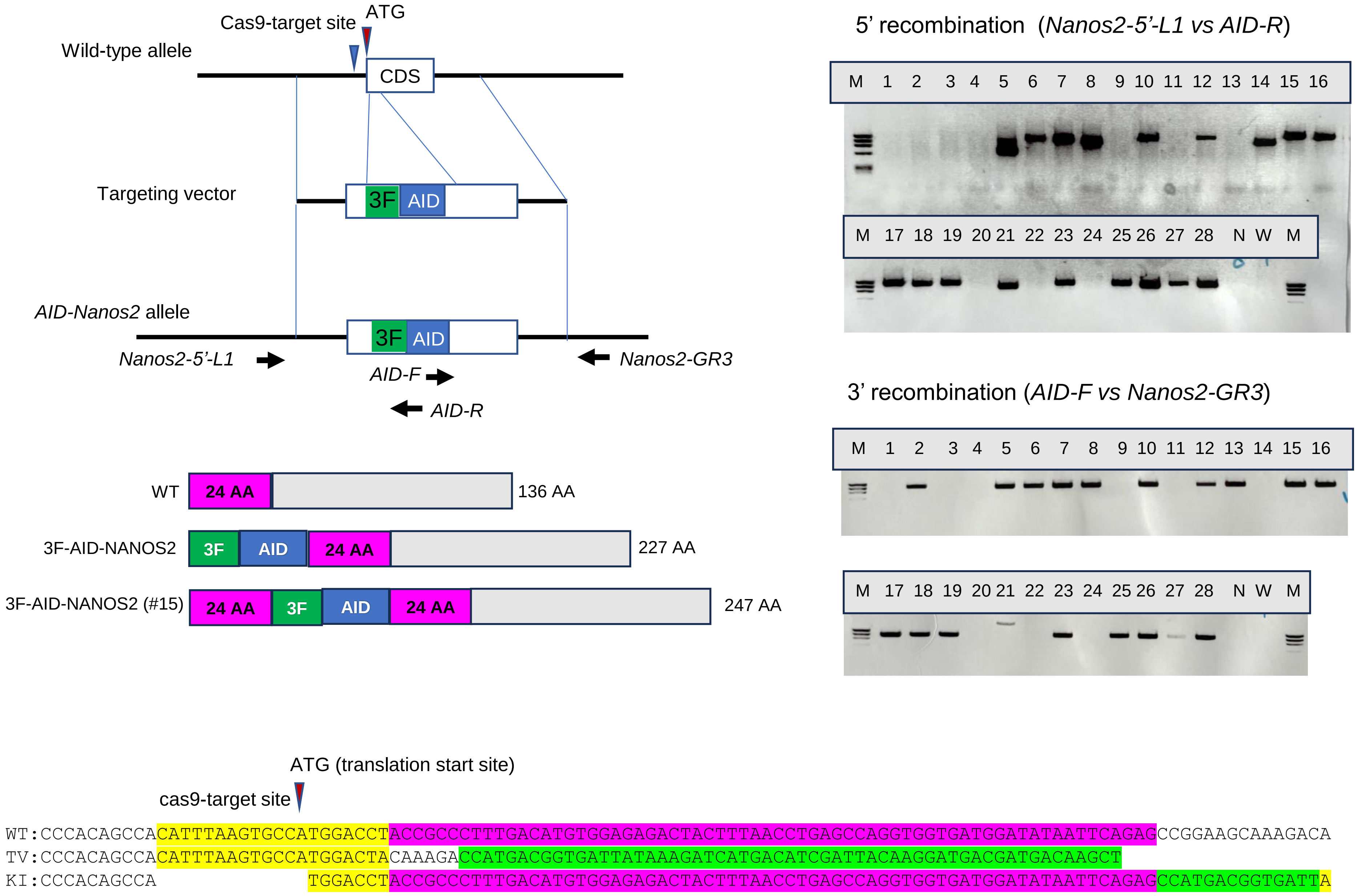
Cas9-target site
ATG
Wild-type allele
CDS
Targeting vector
3F
AID
AID-Nanos2 allele
3F
AID
Nanos2-5'-L1
AID-F
AID-R
Nanos2-GR3
5' recombination (Nanos2-5'-L1 vs AID-R)
M 1 2 3 4 5 6 7 8 9 10 11 12 13 14 15 16
M 17 18 19 20 21 22 23 24 25 26 27 28 N W M
3' recombination (AID-F vs Nanos2-GR3)
M 1 2 3 4 5 6 7 8 9 10 11 12 13 14 15 16
M 17 18 19 20 21 22 23 24 25 26 27 28 N W M
WT
24 AA
136 AA
3F-AID-NANOS2
3F
AID
24 AA
227 AA
3F-AID-NANOS2 (#15)
24 AA
3F
AID
24 AA
247 AA
ATG (translation start site)
cas9-target site
WT:CCCACAGCCACATTTAAGTGCCATGGACCTACCGCCCTTTGACATGTGGAGAGACTACTTTAACCTGAGCCAGGTGGTGATGGATATAATTCAGAGCCGGAAGCAAAGACA
TV:CCCACAGCCACATTTAAGTGCCATGGACTACAAAGACCATGACGGTGATTATAAAGATCATGACATCGATTACAAGGATGACGATGACAAGCT
KI:CCCACAGCCA TGGACCTACCGCCCTTTGACATGTGGAGAGACTACTTTAACCTGAGCCAGGTGGTGATGGATATAATTCAGAGCCATGACGGTGATTA

Fig. S2

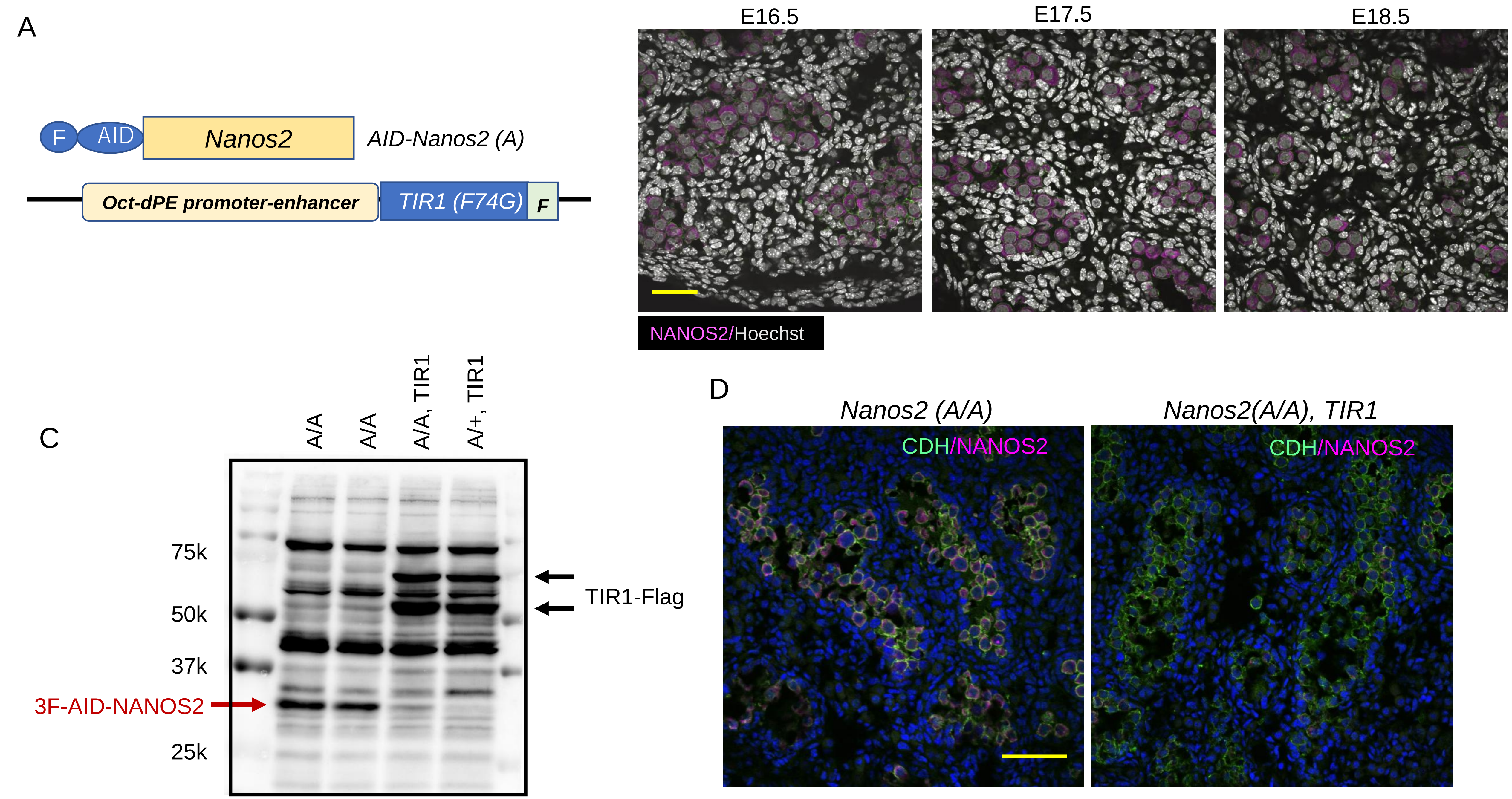
A
F
AID
Nanos2
AID-Nanos2 (A)
Oct-dPE promoter-enhancer
TIR1 (F74G)
F
B
E16.5
E17.5
E18.5
NANOS2/Hoechst
C
A/A
A/A
A/A, TIR1
A/+, TIR1
75k
50k
37k
25k
3F-AID-NANOS2
TIR1-Flag
D
Nanos2 (A/A)
Nanos2(A/A), TIR1
CDH/NANOS2
CDH/NANOS2

Fig. S3

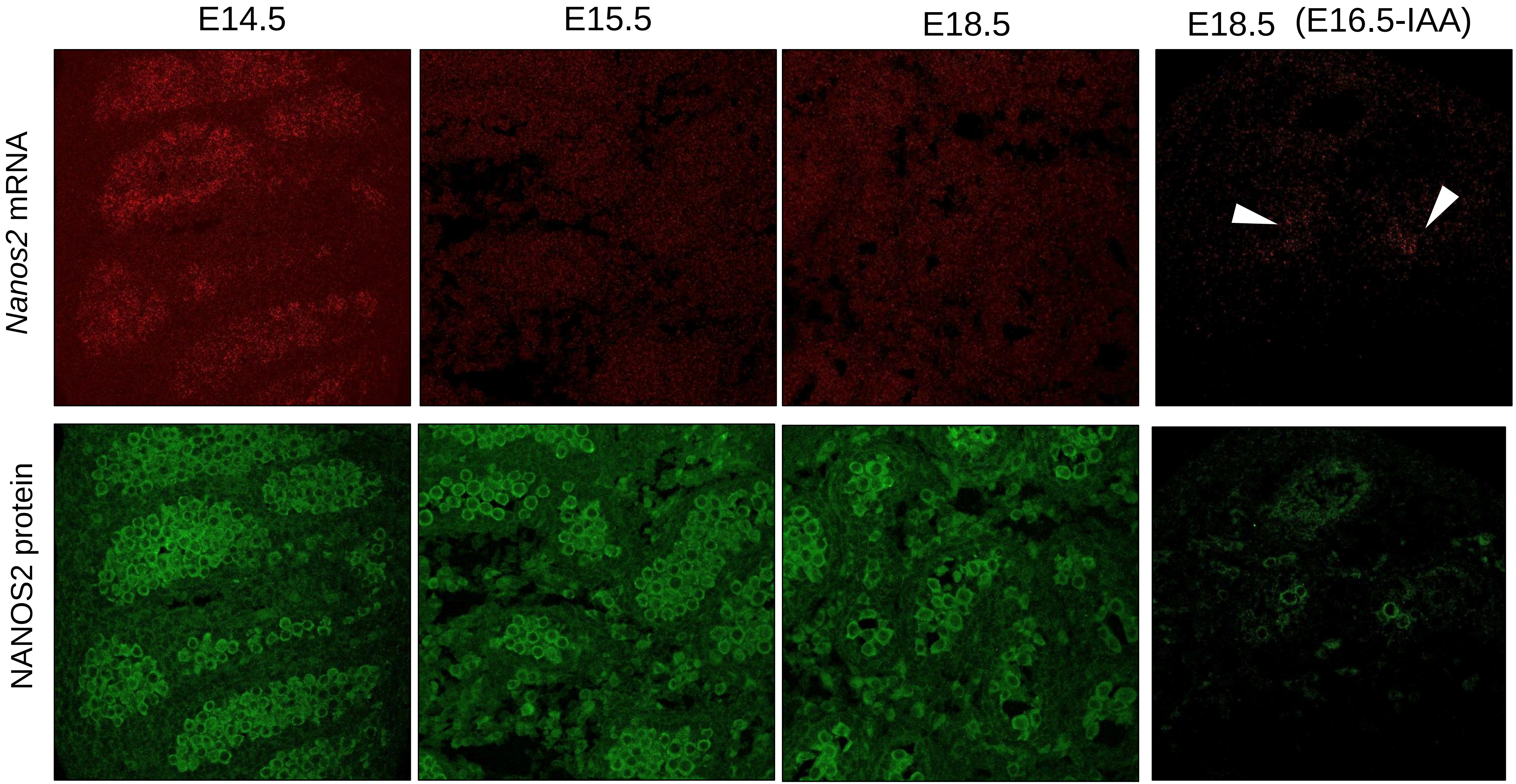

Fig. S4

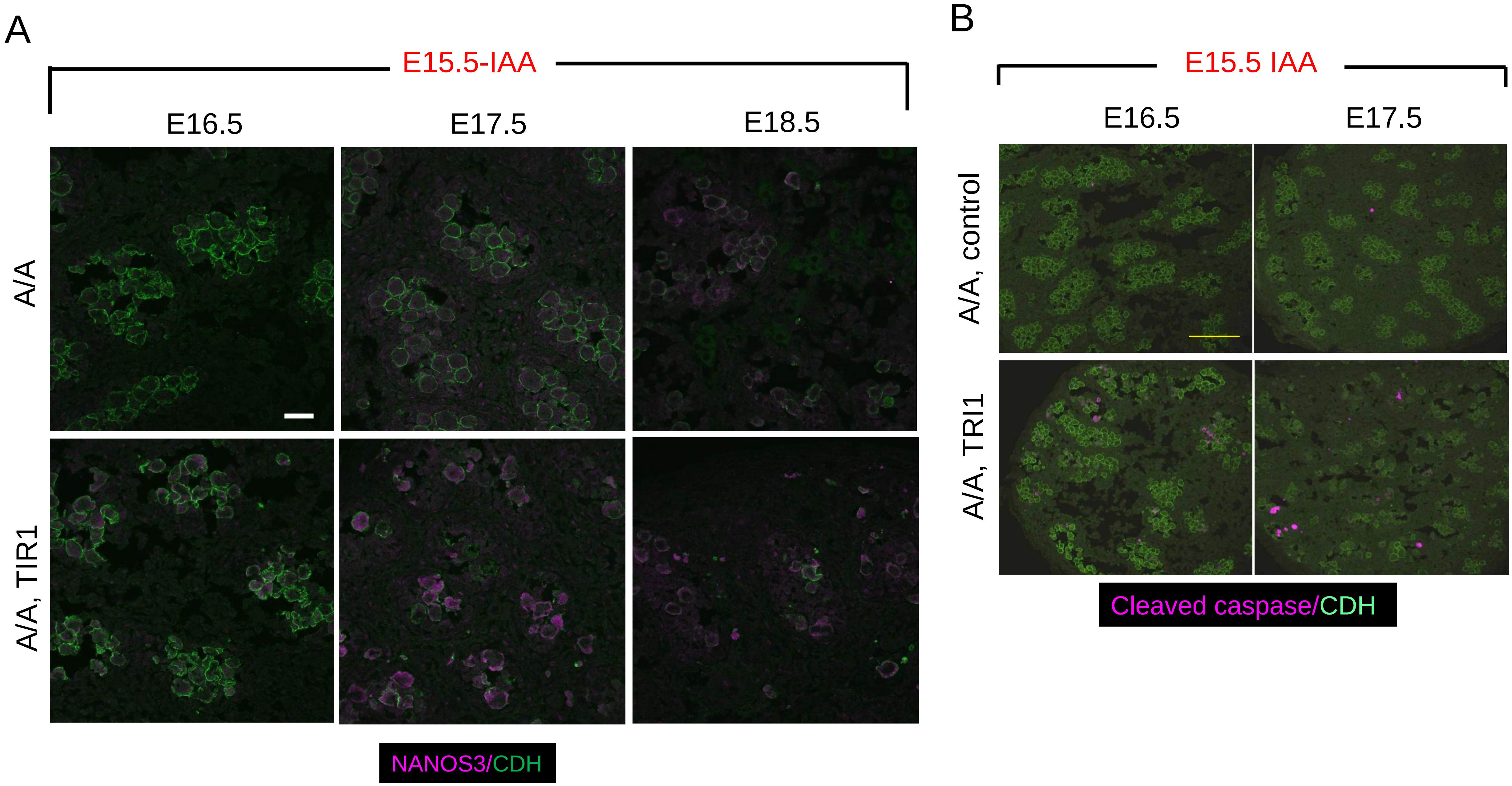
A
E15.5-IAA
E16.5
E17.5
E18.5
A/A
A/A, TIR1
NANOS3/CDH
B
E15.5 IAA
E16.5
E17.5
A/A, control
A/A, TRI1
Cleaved caspase/CDH

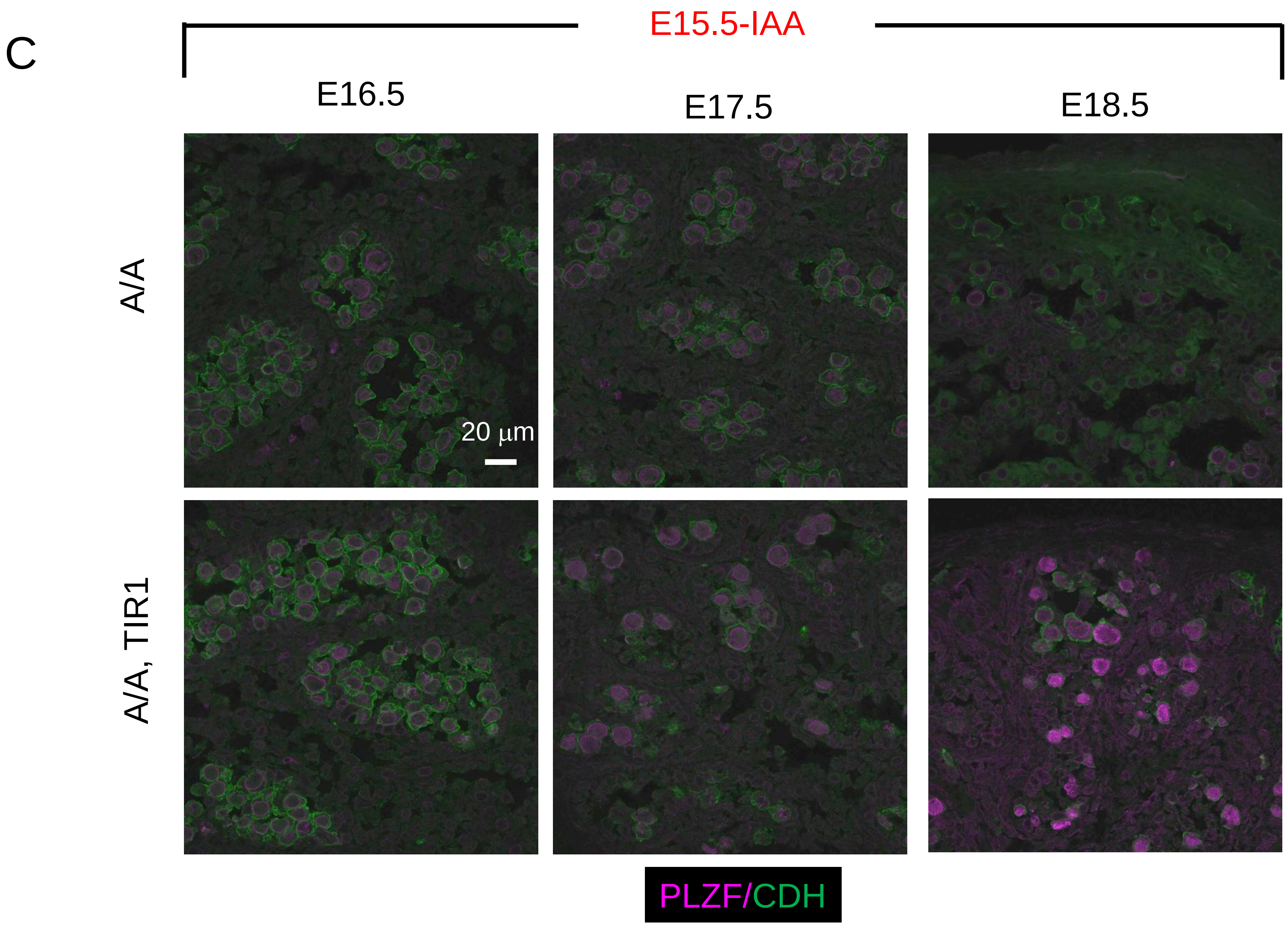
C
E15.5-IAA
E16.5
E17.5
E18.5
A/A
A/A, TIR1
20 µm
PLZF/CDH

Fig. S5

### Summary of Data

| Time of IAA treatment/sampling stage | | | | | | NANOS2 | DNMT3L | Ki67 | Caspase | NANOS3 |
|---|---|---|---|---|---|---|---|---|---|---|
| | | E15.5 | E16.5 | | | - | + | - | + | + |
| | | E15.5 | | E17.5 | | - | +/- | ++ | ++ | ++ |
| | | E15.5 | | | E18.5 | ++ | +/- | ++ | + | ++ |
| | | | E16.5 | E17.5 | | - | ++ | - | + | +/- |
| | | | E16.5 | | E18.5 | ++ | ++ | + | + | ++ |
| | | | | E17.5 | E18.5 | - | ++ | - | + | - |
| Control | | E15.5 | | | E18.5 | ++ | ++ | - | - | - |

### Table S1: Actual number of GFRA1-positive cells among PLZF-positive undifferentiated spermatogonia

| Stage | Sample | PLZF(GFRA1) Number of cells / 0.4 $mm^2$ | | | | |
|---|---|---|---|---|---|---|
| 1M | Control | 48(4) | 31(4) | 22(1) | 29(4) | |
| | E15.5+E16.5-IAA | 75(6) | 78(10) | 61(10) | 82(12) | |
| | E16.5-IAA | 97(11) | 98(17) | 83(8) | 15(0) | |
| 2.5M | control | 46(3) | 25(4) | 29(2) | | |
| | E16.5-IAA | 67(5) | 51(5) | 43(5) | 22(1) | 36(6) |
| 3.75M | control | 62(3) | 56(4) | | | |
| | E17.5-IAA | 42(5) | 47(1) | | | |
| | E18.5-IAA | 31(6) | 31(0) | 39(5) | | |
| 6M | control | 59(5) | 39(3) | 45(2) | | |
| | E16.5-IAA | 4(0) | 4(2) | | | |
